# Understanding the development of enzalutamide resistance based on a functional single-cell approach

**DOI:** 10.1101/2024.10.20.619319

**Authors:** Changhui Xue, Hyun-Kyung Ko, Kasen Shi, Janet Pittsenbarger, Lucien Vu Dao, Kaiyo Shi, Maximilian Libmann, Hao Geng, David Z. Qian

**Affiliations:** Division of Oncological Sciences Knight Cancer Institute Oregon Health & Science University Portland, Oregon USA

## Abstract

Most metastatic prostate cancers (PCa) initially depend on androgen for survival and proliferation. Thus, anti-androgen or castration therapies are the mainstay treatment. Although effective at first, androgen-dependent PCa (ADPC) universally develops therapy resistance, thereby evolving to the incurable disease, called castration resistant PCa (CRPC). Currently, mechanisms underlying the emergence of CRPC from ADPC are largely unclear. We used single-cell RNA-sequencing (scRNA-Seq) to determine how a therapy-naïve ADPC cell line – LNCaP responds to the anti-androgen drug, enzalutamide. We found that most cells expressed the drug-target androgen receptor (AR+), while a small subpopulation (∼12%) expressed low or no AR (AR^low/-^). Gene set enrichment analysis (GSEA) revealed that AR+ and AR^low/-^ cells were enriched with significantly different gene expressions and signaling pathways. Unexpectedly, AR^low/-^ cells displayed robust transcriptional response, including upregulations of genes and pathways involved in clinical CRPC. Next, we isolate AR^low/-^ and AR+ cells from the LNCaP cell line, and functionally confirmed the enzalutamide resistant phenotype of AR^low/-^ cells in vitro and in xenograft models in vivo. Finally, to explore a therapeutic option for AR^low/-^ cells, we found that AR^low/-^ cells expressed low levels of NAD+ biosynthesis genes, notably NAPRT, indicating a possible vulnerability to inhibitors blocking NAD+ synthesis. Indeed, treating AR^low/-^ cells with NAD+ synthesis inhibitors, FK866 and OT-82, significantly inhibited the survival and proliferation of AR^low/-^ cells, thus suggesting a possible novel therapeutic option for ADT and enzalutamide resistant PCa.

**SUMMARY:** Single-cell RNA-Sequencing reveals heterogeneities of tumor cell populations. In most cases, however, the functional significance of the observed heterogeneity is not tested. In this study, we first identified a possible therapy-resistant prostate cancer cell subpopulation with scRNA-Seq, then confirmed the resistant phenotype with single cell and colony – based cloning and functional testing. In addition, we also identified a therapeutic vulnerability of the resistant cells.

## INTRODUCTION

Metastatic prostate cancers are primarily treated by therapies blocking the androgen/androgen receptor (AR) axis^1^. This is because prostate cancer cells are initially androgen (male steroid sex hormone) dependent (ADPC). Androgen activates AR by binding to AR and promoting its nuclear localization and dimerization. In turn, AR becomes a master transcription regulator and activates gene expressions essential for tumor cell survival and proliferation^2^. Clinical anti-androgen/AR treatments include castration or androgen-deprivation (ADT) or in the past 10 years enzalutamide (Enz)^1, 3^. Although initially effective, the long-term efficacy and durability of these treatments are limited by the development of resistance ^2, 4^. As a result, the cancers progress to a stage called CRPC (castration resistant prostate cancer), which currently is incurable and causing 27,000 to 30,000 deaths per year in the US.

Multiple AR dependent and independent molecular pathways have been implicated in the development of ADT/Enz resistance ^2, 5^. In the AR-dependent setting, AR activation (nuclear localization and DNA binding) becomes independent of androgen ^6^. Further, the transcriptional activity of AR is reprogrammed to regulate more gene expressions, many of which may support CRPC ^6^. In the AR-independent setting, additional oncogenic pathways, such as Myc, PI3K/AKT, E2F, HIF1, GR and EMT, are activated to play more important roles in supporting tumor cells^2^. At the cellular level, AR-expressing (AR+) tumor cells may undergo neuroendocrine differentiation (NED), while losing AR expression, thereby becoming resistant to androgen/AR-targeted therapies ^7^. Despite these understanding, there are still significant knowledge gaps. It is largely unclear how AR becomes androgen independent, how AR-independent oncogenic pathways are activated, and what are the vulnerabilities of ADT/Enz-resistant tumor cells.

Tumor heterogeneity plays a significant role in promoting treatment resistance^8–10^. Prostate cancers are known to be heterogenic^11^. In this study. we hypothesized that the heterogeneity of prostate cancers may manifest at the therapy response and resistant levels, with subsets of cells are less responsive to the treatment, and therefore clonally selected and expanded to confer resistance.

## RESULTS

### LNCaP cell line has gene/pathway heterogeneities, especially androgen/AR signaling

As previously described ^12^, we cultured the AR-expressing LNCaP cells in androgen-stimulated condition, and transiently treated it with anti-androgen enzalutamide (Enz) or solvent control. Afterwards, we used single-cell RNA-Seq to determine potential heterogeneities of transcriptome and Enz-response. In both conditions, unsupervised-high dimensional clustering (UMAP) revealed 4 distinctive cell clusters (Figure 1A). In the control condition, differential gene expression analysis revealed that cluster 2 (c2) and 3 (c3), which represented ∼12% of the total population (Figure 1B), had very different gene expression pattern from cluster 0 (c0) and 1 (c1) (Figure 1C). Notably, cells in c2/c3 expressed none/low levels of AR and canonical androgen/AR target genes (Figures 1D-1F).

**Figure 1.**
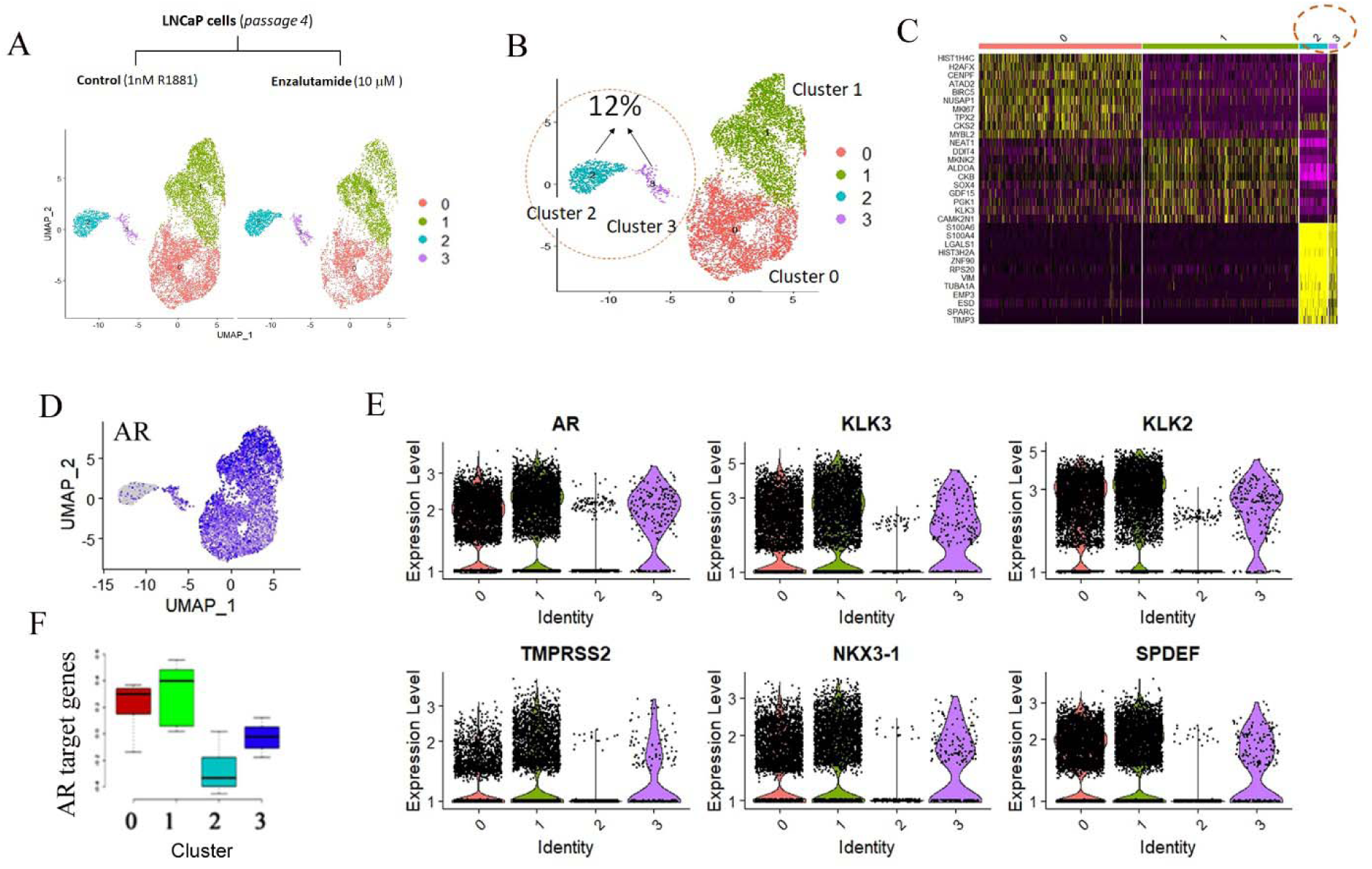
Single-cell RNA-Seq reveals that LNCaP cell line has subpopulations that are low in AR expression and signaling (AR-low). **(A)** scRNA-Seq of early passage LNCaP cells in response to androgen (1 nM R1881) stimulated control or 10 μM enzalutamide (Enz) for 24 hours. The sequencing data was analyzed with UMAP in Seurat R packages. **(B – F)** In the control condition, LNCaP cells have subpopulations, clusters (c) 0 – 3, in which clusters 2 and 3 have low expressions for AR and AR-target genes. (B) The % of cells (clusters 2 and 3) with low expressions for AR and AR-target genes. (C) Heatmap of differentially-expressed genes (DEG) in LNCaP subpopulations (clusters 0 – 3). (D) Feature plot of AR-expressing cells (blue). (E) Violin plots of AR-target genes in subpopulations (clusters 0 – 3). (F) Box and whisker plot of AR-target gene expressions in 4 subpopulations. The AR-target genes are defined by Molecular Signature Database (MSigDB).

We next labelled the cells in c2/c3 as AR^low/-^ and the c0/c1 counterparts as AR^+^, respectively. In contrast to low AR signaling, AR^low/-^ cells were enriched with genes of the epithelial-mesenchymal transition (EMT) pathway (Figures 2A-2C), and genes with oncogenic activities, some of which may contribute to CRPC and Enz-resistant ^2^, e.g. glucocorticoid receptor (GR) – NR3C1 (Figure 2D). By mining the signature genes in AR^low/-^ cells with bulk RNA-Seq dataset of CRPC patients from the SU2C/PCF dream team study, we further found that some of the c2/c3 cell signatures (Figure 2E) were significantly associated with poor overall survival of CRPC patients (Figure 2E). The differential gene expression between AR^+^ (c0/c1) and AR^low/-^ (c2/c3) subpopulations can also be reflected at the pathway level (Figures 2F – 2H). Notably, AR^low/-^ cells were enriched with pathways involved in CRPC, e.g. hypoxia (Figure 2F), extracellular matrix (Figure 2G), and stem cells (Figure 2H).

**Figure 2.**
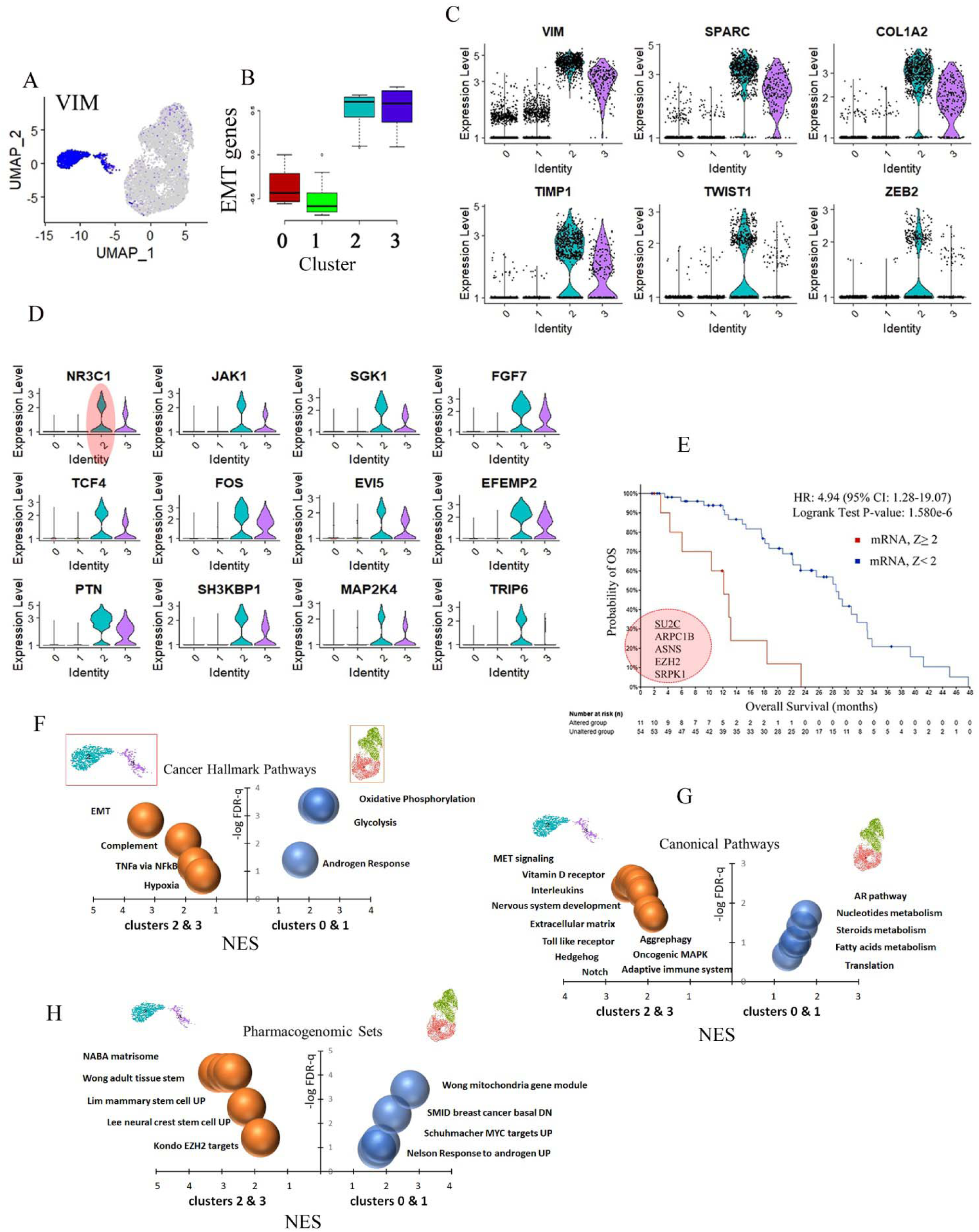
The AR-low (AR^low/-^) subpopulation is enriched with genes and pathways that are distinctively different from the AR-high (AR^+^) cells. **(A-C)** AR^low/-^ cells (clusters 2/3, c2c3) are enriched with EMT genes. (A) Feature plot of cells expressing EMT marker VIM (blue). (B) Box whisker plot of EMT target gene expressions in subpopulations. The EMT target genes are defined by MsigDB. (C) Violin plots of representative EMT genes in subpopulations (clusters 0 – 3). **(D)** Violin plots of selective genes, many of which are known to be involved in prostate cancer and enzalutamide resistance. **(E-F)** Selective signature genes of cluster 2 and 3 are significantly associated with poor overall survival of CRPC patients. (E) Cells with expressions of *ARPC1B*, *ASNS*, *EZH2*, and *SRPK1* were visualized in UMAP. (F) Overall survival analysis of selective AR^low/-^ signature genes in (E) using bulk RNA-seq dataset from SU2C/PCF dream team. **(G – I)** GSEA reveals that AR^low/-^ and AR^+^ cells are enriched with genes for distinctively different pathways for cancer hallmark (G), canonical signaling pathways (H), and chemical and genetic perturbations (I).

### LNCaP displays heterogenic response to enzalutamide

We next determined how AR^low/-^ and AR^+^ cells responded to enzalutamide. As expected, the AR^+^ cells responded to Enz-treatment with mostly downregulation of gene expressions (Figure 3A), many of which are canonical target genes for androgen/AR-signaling, e.g. PSA, KLK2, and NKX3-1 (Figure 3A). Interestingly, AR^low/-^ cells responded to Enz-treatment with both downregulation and upregulation of gene expressions (Figures 3A – 3B). Some of the Enz-upregulated genes are known to be involved in anti-androgen resistance ^13–15^, e.g. BRD2 and TIMP1 (Figure 3B). A subset of these genes (NAXE, FTL, HMGN2, RRM2, ID3, S100A10, TUBA1A, TUBB2A) is significantly upregulated (Figure 3C), and is associated with poor overall patient survival in clinical CRPC (Figure 3D). Further, GSEA revealed that AR^low/-^ subpopulation responded to Enz-treatment with the upregulation of different pathways from the AR^+^ cells (Figure 3E), some of which are known to mediate castration resistance ^16–18^, e.g. autophagy and E2F.

**Figure 3.**
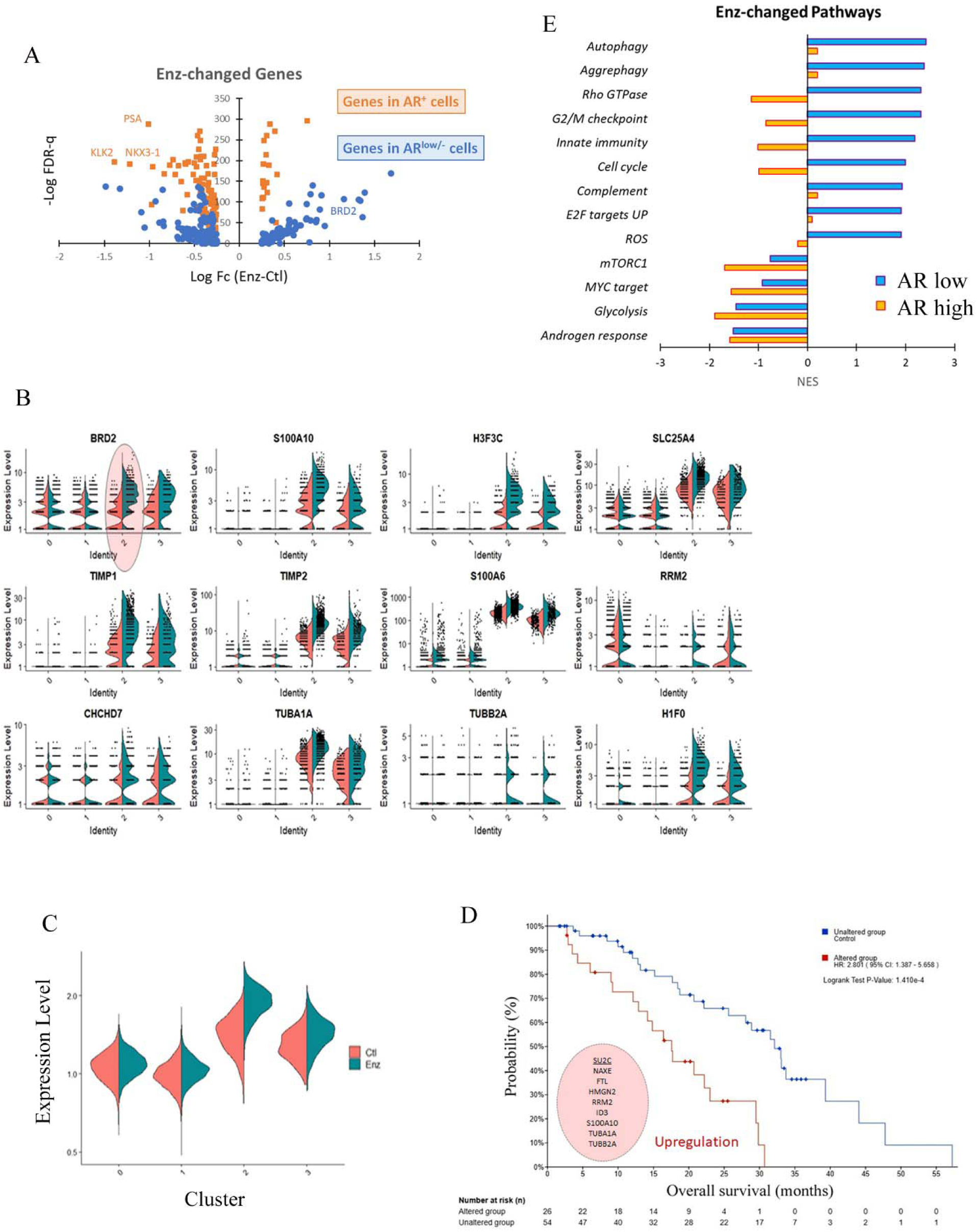
AR^low/-^ and AR^+^ cells display differential response to enzalutamide. **(A)** Volcano plot of genes in response to Enz in AR^low/-^ (clusters 2 – 3) and AR^+^ (clusters 0 – 1) cells. Some of the genes that were downregulated by Enz in AR^+^ cells (PSA, KLK2, NKX3-1) or upregulated by Enz in AR^low/-^ cells (BRD2) are labelled. **(B)** Violin plots of selected genes that are upregulated in AR^low/-^ cells (clusters 2 and 3) in response to Enz. Split violin is used to represent control (orange color) or Enz-treated (green) condition. **(C)** A selective group of genes (*NAXE*, *FTL*, *HMGN2*, *RRM2*, *ID3*, *S100A10*, *TUBA1A*, *TUBB2A*) in c2 and c3 clusters is upregulated by Enz. Top: feature plot, bottom: violin plot. (D) The upregulation of geneset in (C) is significantly associated with poor overall survival in CRPC patients, using bulk RNA-seq dataset from SU2C/PCF dream team. **(E)** Pathways that are changed in response to Enz in AR^low/-^ and AR^+^ cells.

### Isolation of the AR^low/-^ cells

Similar to the emergence of therapy-resistance in the clinics, therapy-sensitive LNCaP cell line is known to become resistant following chronic anti-androgen treatments in vitro or in vivo ^19–22^. However, the underlying mechanisms are not fully understood. In the LNCaP cell line, a small PSA-negative or low (PSA^−/low^) subpopulation was previously reported to cause castration resistance after long-term castration culture ^23^. The identification of this population was based on the transfection of PSA-promoter-driven GFP plasmid ^24^. Based on our single-cell results above, we hypothesized that the AR^low/-^ subpopulation may also represent an enzalutamide-resistant lineage. To test this, we set to isolate the AR^low/-^ cells from the parental LNCaP cell line. As shown in Figure 2H and Figure 4A, the AR^low/-^ cells were enriched with genes along the stem and cancer stem-like pathways, which are known to facilitate the formation of cell colonies (Figure 4A) ^24, 25^. Thus, we plated parental LNCaP cells in single-cell suspension to allow the formation of single cell-derived colonies, and transferred individual colonies to 96-well plate to continue cell culture in the presence of androgen (1 nM R1881) (Figure 4B). Over the course of 3 months, we established ∼50 single cell/colony derived LNCaP clones. Next, we used qRT-PCR to determine the levels of AR and EMT marker vimentin (VIM) in individual clones (Figure 4C), and identified 3 clones with AR^low/-^ and AR^+^ expression phenotypes, respectively (Figures 4C – 4D).

**Figure 4.**
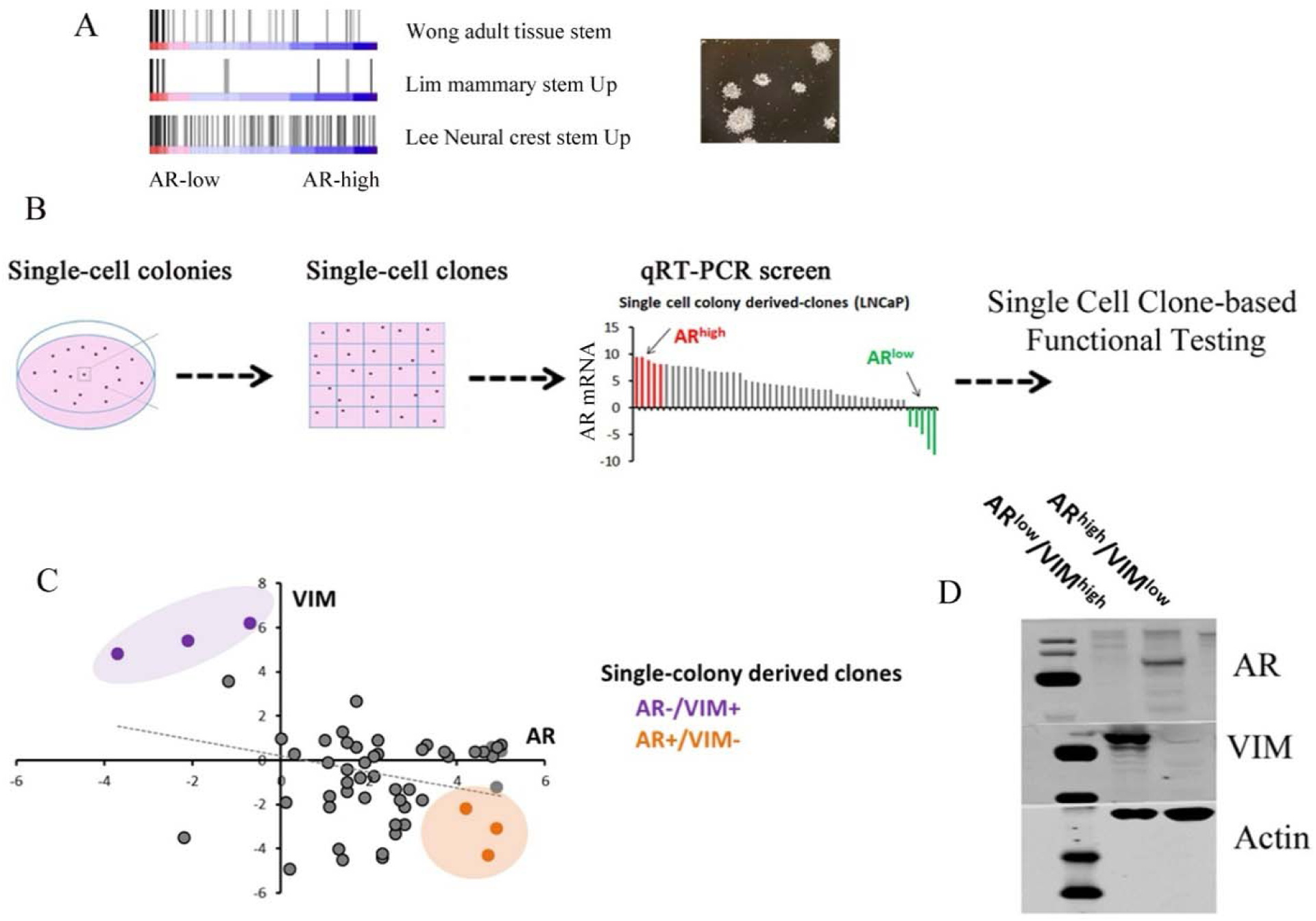
Isolation of the AR^low/-^ subpopulations. **(A)** GSEA reveals that AR^low/-^ cells were enriched with stem cell pathway genes, and example of colony formation, in which 500 early passage LNCaP cells in single-cell suspension were plated with R1881-control media to form single cell – derived colonies over 7 – 10 days. **(B)** Schematics of isolation of AR^low/-^ subpopulation and downstream functional testing. **(C)** Results of qRT-PCR of AR and VIM from ∼50 single-cell/colony-derived LNCaP clones, using schematics in (B). The x and y axis are mRNA levels of AR and VIM in relationship to actin in individual clones. The clones in shaded areas were identified as AR^low/-^ and VIM-high (purple color) or AR^+^ and VIM-low (orange color). **(D)** Representative of western blots of AR, VIM and beta-actin (as loading control) in qRT-PCR identified AR^low/-^ and AR^+^ cells.

### Functional validation of Enzalutamide-resistance by the AR^low/-^ clone

We treated the isolated AR-high or AR-low clones with ADT or ADT plus Enz. We found that the AR^low/-^ clone was indeed insensitive or resistant to the treatment, compared to the AR^+^ counterparts (Figures 5A-5B). Next, we stably transfected RFP and GFP expression plasmids into AR^low/-^ and AR^+^ cells, respectively, and plated the RFP+/AR^low/-^ and GFP+/AR^+^ cells at 1:1 ratio (Figure 5C). As expected, in ADT/Enz conditions, the GFP+/AR^+^ cells were significantly inhibited, while the RFP+ AR^low/-^ cells were resistant and proliferated (Figures 5D, 5G). On the other hand, the GFP+/AR^+^ cells displayed better proliferation pattern in R1881-condition (Figures 5E, 5H), unless the media was switched to ADT/Enz (Figures 5F, 5H). In vivo, we established subcutaneous xenografts with similar amount of AR^+^, AR^low/-^, or 4:1 (AR^+^: AR^low/-^) mixture cells. The tumor bearing mice were treated with Enz as previously described ^12^. We found that the AR^+^ tumors were sensitive to enzalutamide (Figure 5I), while AR^low/-^ and 9:1 (AR^+^: AR^low/-^) mixture tumors were insensitive / resistant (Figures 5J – 5K).

**Figure 5.**
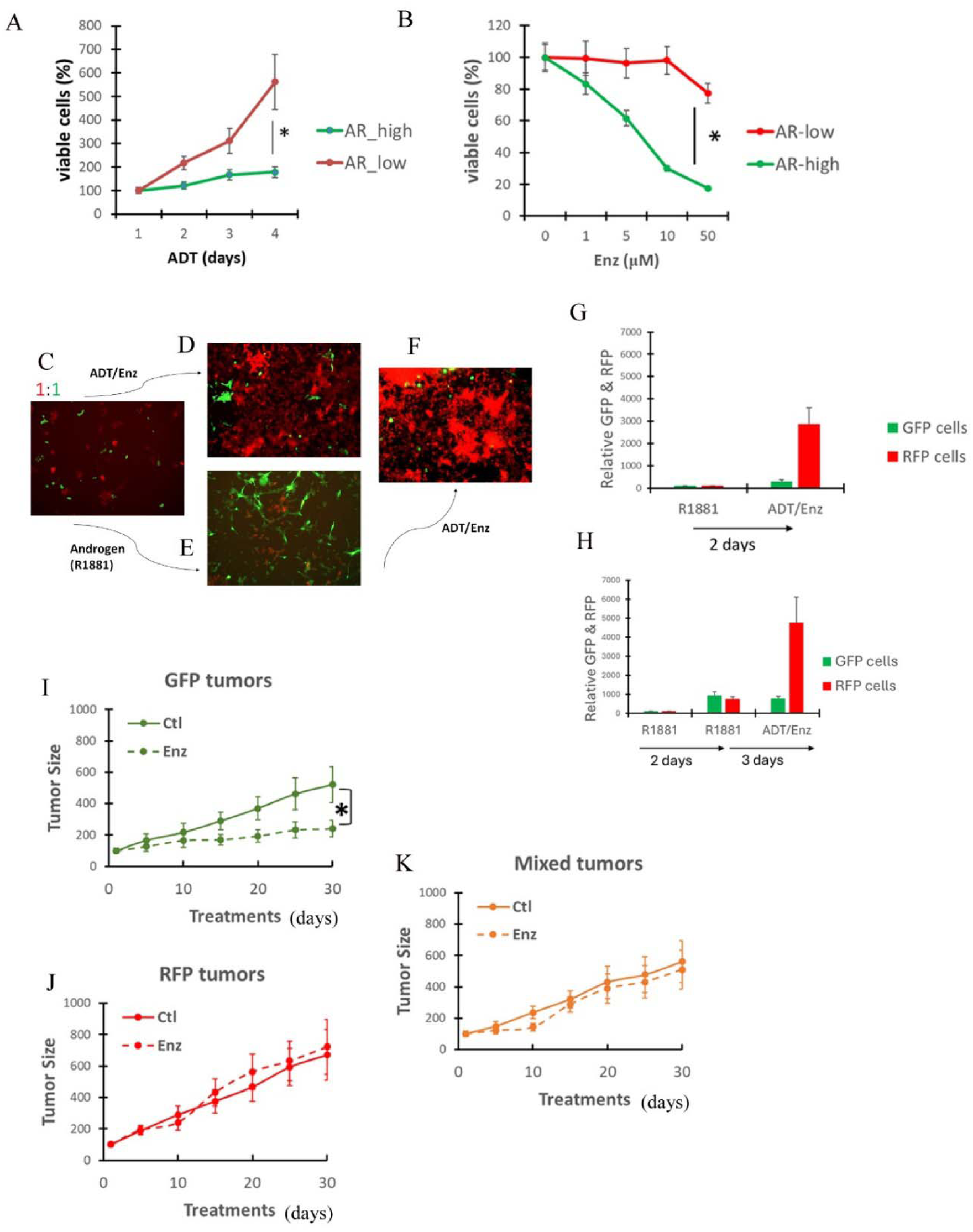
Functional validation of AR^low/-^ cells in vitro and in vivo in response to ADT and enzalutamide. **(A-B)** The viability and proliferation of AR^+^ and AR^low/-^ cells in response to ADT with 5% charcoal-stripped serum (CSS) in (A) or ADT plus increasing concentrations of Enz in (B). **(C – F)** Representative images of GFP+/AR^+^ and RFP+/AR^low/-^ cells, 24 hours after 1:1 plating in 5% CSS + 1 nM R1881 media (C), followed by 48 hours in ADT/Enz (5% CSS/10 μM) (D), or 48 hours in 1 nM R1881 media (E) and switched to ADT/Enz for additional 72 hours (F). **(G – H)** Relative quantification of GFP and RFP intensity for cells in D – F. **(I – K)** Xenografts of AR^+^ (I), AR^low/-^ (J), or 9:1 mix of AR^+^ and AR^low/-^ (K) tumors in response to vehicle control or Enz. All * indicates P < 0.05, t-test, n = 3 (in vitro) or = 6 (in xenograft).

### The AR^low/-^ clone is vulnerable to NAD+ synthesis inhibitors

To explore a therapeutic approach for the ADT/Enz-resistant clone (* in Figure 6A), we treated cells with a panel of inhibitors targeting, aurora A kinase (Alisertib)^26^, BET (JQ1)^27^, EZH2 (Tazemetostat)^28^, GR (RU486)^29^, PARP (Olaparib)^30^ and EMT (FiVe1)^31^, all of which are being tested in clinical or preclinical settings against CRPC. However, none elicited significantly different response (sensitivity) between AR^+^ and AR^low/-^ cells (Figure 6A). Next, we used bulk RNA-seq to compare the transcriptomes of AR^low/-^ and AR^+^ cells in depth, and found a more extensive list of genes that were differentially expressed between AR^low/-^ and AR^+^ clones (Figure 6B). Gene Ontology (GO) analysis indicted that AR^low/-^ cells were enriched with the downregulation of genes mediating NAD+ biosynthetic pathways (Figure 6C)^32^. NAD+ is an essential co-factor for cell energy metabolism, oxidative stress response, and chromatin modification ^33^. It can be synthesized via 3 pathways ^32^, in which NAMPT and NAPRT are rate limiting enzymes ^33^. We found that AR^low/-^ and AR^+^ cells expressed equivalent levels of NAMPT, while the NAPRT mRNA and protein levels were significantly lower in AR^low/-^ cells (Figure 6D – 6E). Further, the cellular NAD+ level in AR^low/-^ cells was also significantly lower than AR^+^ cells (Figure 6F).

**Figure 6.**
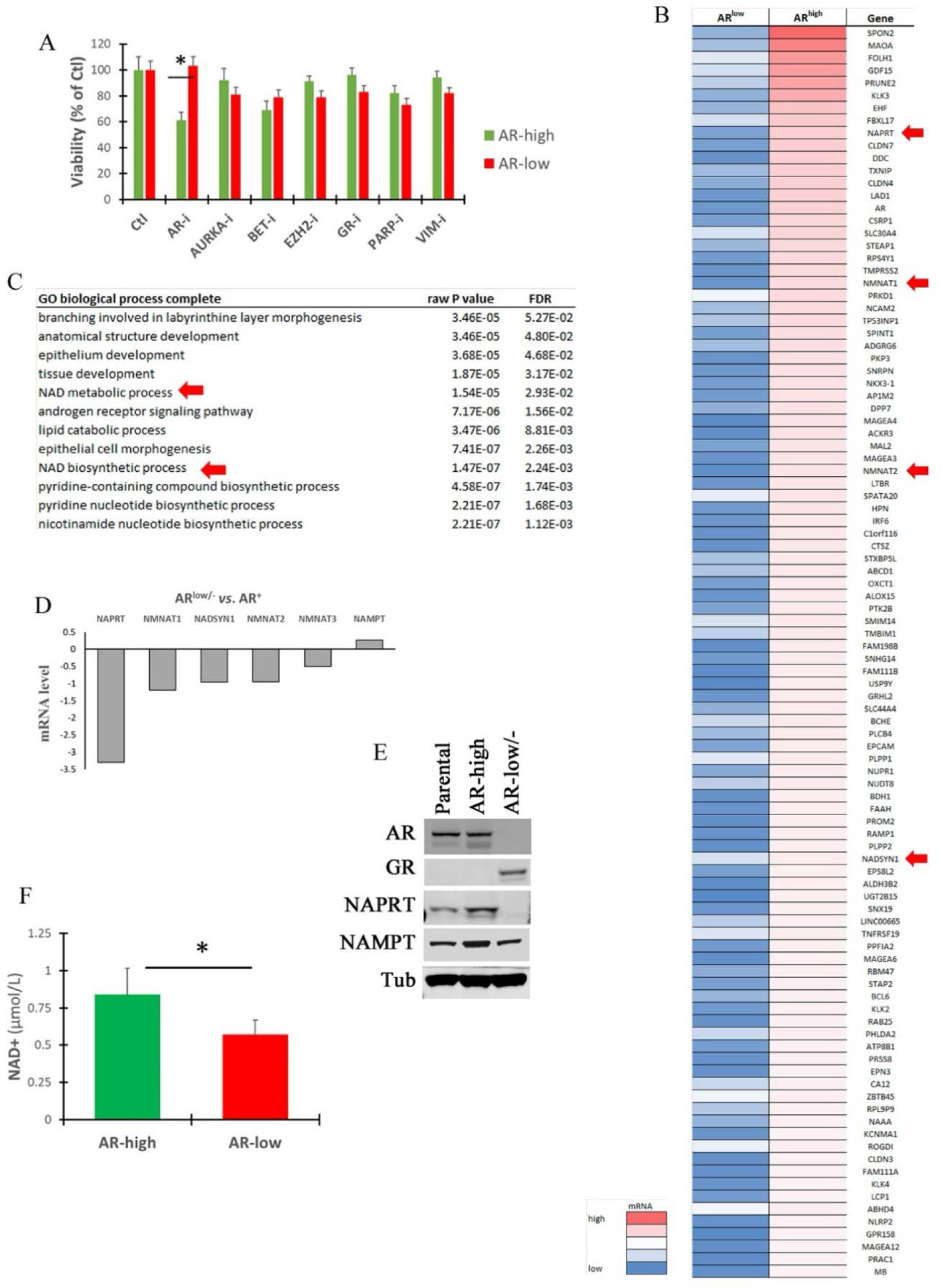
AR^low/-^ cells have lower expression of NAPRT than AR^+^ cells. **(A)** AR^low/-^ and AR+ cells were treated by a panel of inhibitors targeting androgen/AR (AR-i, 10 µM Enz), aurora A kinase (100 nM Alisertib), BET (250 nM JQ1), EZH2 (5 µM Tazemetostat), GR (5 µM RU486), PARP (5 µM Olaparib), and EMT/Vimentin (1 µM FiVe1). The doses used were based on literatures and considered to be physiologically relevant. * Indicates P < 0.01, t-test. **(B)** Genes that were down-regulated in AR^low/-^ as compared to AR+ cells based on bulk RNA-Seq. Red arrow indicates genes mediating NAD+ biosynthesis. **(C)** GO analysis of genes that were significantly downregulated in AR^low/-^ cells comparing to AR+ cells. Red arrow indicates the NAD+ biosynthesis pathways. **(D)** Bulk RNA-Seq results of genes involved in NAD+ biosynthesis in AR^low/-^ and AR+ cells. **(E)** Representative western blots of NAPRT and NAMPT protein expressions, and NAD levels in AR^low/-^ *vs*. AR+ cells. **(F)** NAD+ levels in AR^low/-^ and AR+ cells. * or # Indicate P < 0.05, n=3, t-test.

In the literature, tumor cells lacking NAPRT are reported to have enhanced sensitivity to inhibitors blocking NAMPT ^34–36^. Thus, we treated AR^+^ and AR^low/-^ clones with two NAMPT inhibitors, FK866 and OT-82 ^36, 37^. Notably, AR^low/-^ cells, but not the AR^+^ counterparts, displayed a dose-dependent sensitivity (Figure 7A). In parallel experiments, we found cellular NAD+ levels were significantly reduced by the inhibitors in AR^low/-^ cells, but to much less extent in AR^+^ counterparts (Figure 7B). When we added NAMPT-downstream product nicotinamide mononucleotide (NMN) into the culture media of AR-low cells, the cellular NAD+ level was restored (Figure 7C), and cell viability was also rescued (Figure 7D). Next, we reduced NAPRT in AR^+^ cells by siRNA, which significantly increased the sensitivity to NAMPT inhibitors (Figure 7E). In contrast, a plasmid-based overexpression of NAPRT in AR^low/-^ cells reduced the NAMPT inhibitor sensitivity (Figure 7F). To determine whether the differential sensitivity can be recapitulated in other prostate cancer cell models, we determine the NAPRT and NAMPT levels in additional ADT-resistant prostate cancer cells - C42B, 22Rv1, PC3, and DU145 (Figure 7G). We found that PC3 cell had the lowest level of NAPRT, and was the most sensitive to OT-82 (Figure 7H). Further, silencing NAPRT with siRNA increased OT-82 sensitivity in C42B, 22Rv1 and DU145 cells (Figure 7H). Taken together, we find a previously unknown vulnerability of CRPC, in which the low level of NAPRT expression may confer sensitivity to inhibitors targeting NAMPT.

**Figure 7.**
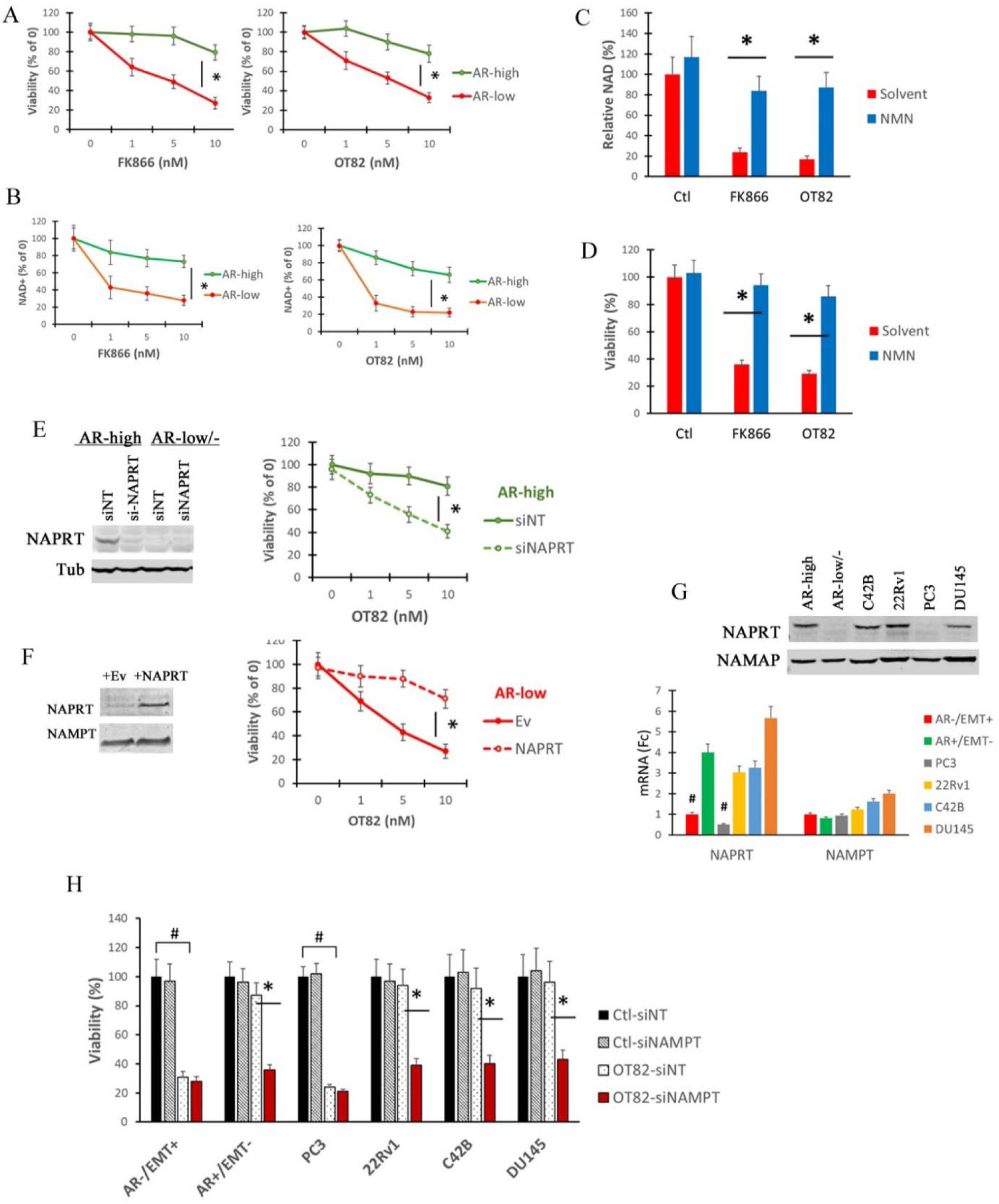
AR^low/-^ cells are vulnerable to NAMPT inhibitors due to the lack of NAPRT. **(A)** The viabilities of AR^low/-^ and AR^+^ cells after being treated by increasing doses of NAMPT inhibitors FK866 and OT82 for 3 days. **(B)** In similar experiments to (A), cells treated by NAMPT inhibitors were used to measure cellular NAD+ levels. **(C – D)** In similar experiments to (A), nicotinamide mononucleotide (NMN, 1 mM), the downstream metabolic product of NAMPT and precursor of NAD+, was added into the cell culture media. Cellular NAD+ levels were determined in (C) and viabilities in the presence of NAMPT inhibitors were determined in (D). **(E)** NAPRT-expressing AR^+^ cells were treated with OT-82 in the presence of siRNA cocktails silencing NAPRT (si-NAPRT) or non-targeting control (NT). Afterwards, NAPRT protein levels (left) and cell viabilities (right) were determined. The NAPRT-deficient / AR^low/-^ cell was used as a positive control for siRNA. **(F)** NAPRT-deficient AR^low/-^ cells were transfected with plasmids coding for NAPRT overexpression vector or empty vector (Ev) control and treated with OT82. Afterwards, NAPRT protein levels (left) and cell viabilities (right) were determined. **(G)** NAPRT and NAMPT transcript levels were determined by western blots (top) and qRT-PCR (bottom) in a panel of ADT/Enz-insensitive/resistant PCa cell lines. **(H)** The PCa cell lines in (G) were treated by OT-82 in the presence of siRNA cocktails silencing NAPRT or NT-controls. Cell viabilities in relationship to siNT-treated solvent control are shown. All * indicates P < 0.05, t-test, n = 3.

## Discussion

Single-cell RNA sequencing has been widely used to determine tumor heterogeneity. In most studies, however, the functional significance of the heterogeneity is not confirmed with bench-based investigation. Resistance to ADT and the next-generation anti-androgen, e.g. enzalutamide is a major obstacle in metastatic prostate cancer treatment ^2^. At the cellular level, mesenchymal, neuroendocrine, neural, and cancer stem-like cells are thought to be involved ^2^. These cells share a common feature of expressing none or low level of AR, thus are insensitive to the treatments. However, it is unclear whether these heterogenic AR independent cell types emerge via de novo induction or selection from pre-existing clones.

In this study, we used scRNA-Seq to find that the treatment-sensitive LNCaP cell line has intra-cell line heterogeneity. While overwhelming majority of LNCaP cells express AR and AR-target genes, ∼12% express none or low level of AR. This observation is similar to the LNCaP PSA^−/low^ cell identified previously ^24^, when single-cell RNA-Seq was not available. Here, we further found that besides AR and its target genes, AR-high and AR-low cells also harbor very different gene expression programs and pathways. The AR-low cells express genes for GR, EMT, and stemness, all of which have been found to be implicated in treatment resistance and CRPC ^2^. In AR-low cells, EMT is the most significant signature pathway. Interestingly, a recent clinical study also reported that Enz-resistant patient tumor samples were associated with low AR activity but high EMT^14^. Currently, the biogenesis of the AR-low cells remains unclear. Previous (bulk-cell) studies have suggested that AR+ prostate cancer cells, including LNCaP, may undergo de novo trans-differentiation to become AR-negative cells in response to chronic ADT and Enz^19, 22^. However, since we performed the single-cell analysis with early passage-cells that had not been exposed to in vitro anti-androgen/AR agents, it is unlikely that the AR-low cells in our study were induced by the transient/short-term Enz treatment. We also have followed the LNCaP parental cells for ∼40 passages. Bulk qRT-PCR and western blots did not reveal a significant loss of AR. Thus, it is more likely that the AR-low subpopulation is a pre-existing clone.

As expected, our in vitro and xenograft experiments reveal that the AR-low cells are insensitive or resistant to ADT/Enz treatments. An interesting observation is that these cells do indeed respond to ADT/Enz in vitro at the gene expression level. We found that following the treatment, genes and pathways were significantly upregulated in the AR-low cells, but not in AR-high counterparts (Figure 3). Since these cells lack AR, the mechanism for the upregulation is unclear. Because AR-low cells harbor gene expression phenotypes involved in treatment resistance and CRPC, e.g. GR, BRD2, and EMT, we treated the cells with inhibitors targeting the upregulations. However, the therapeutic advantage, compared to the AR-high counterparts, was not obvious. This suggests that the presence of actionable (targetable) gene expression phenotypes is not necessarily translated to vulnerability or treatment sensitivity, and that individual gene/pathway may not be necessary to confirm growth of AR-low tumor cells. This also presents greater obstacle in developing targeted-therapies against AR-low tumors.

To identify genes and pathways critical to the survival and proliferation of AR-low cells, we found that these cells lack the expression of NAPRT, suggesting vulnerability to NAMPT inhibitors due to synthetic lethality based on previous studies (mostly in other types of cancers) ^35, 36^. Our in vitro test revealed that the AR-low cells were indeed sensitive to NAMPT inhibitor with/without ADT/Enz. In contrast, the plasmid-based NAPRT overexpression rescued cells from the growth inhibitory effect of the NAMPT inhibitors. On the other hand, AR-high counterparts were not sensitive the NAMPT inhibition unless we used siRNA to silence NAPRT. To understand the clinical relevance of these observations, we determined the NAPRT and NAMPT expression in clinical prostate cancer datasets. We found that NAPRT and NAMPT transcripts were negatively associated in primary prostate cancers (TCGA, Spearman’s correlation = −0.161, *P*= 3.287e-4) and CRPC (SU2C, Spearman’s correlation = - 0.178, *P*= 9.911e-3) samples, suggesting high-NAMPT tumors can be potentially deficient at NAPRT and thus vulnerable. The clinical application of NAMPT inhibitors warrants further investigation. Currently, it is unclear 1) the NAPRT threshold of inducing vulnerability to NAMPT inhibitors, and 2) whether there are other factors to modify the vulnerability.

Taken together, we use the LNCaP cell line to demonstrate that the ADT/Enz-sensitive cell line is heterogenous on AR expression, in which the AR-low cells have significantly different gene expression phenotypes from the AR-high counterparts, and are insensitive/resistant to anti-androgen and anti-AR therapies. Most current studies of ADT/Enz-resistance are more focused on the de novo changes induced by the therapy, which in turn confer resistance ^38^. Our study suggests that clonal selection can also be involved. In a heterogeneous tumor, the AR-low clones can be candidates for the selection, which may undergo further changes and differentiations to become CRPC.

## METHODS

### Cell lines, cell culture conditions, and key reagents

Cell lines used in this study are human prostate cancer cell lines, LNCaP, C42B, 22Rv1, DU145 and PC3. The C42B cell line was a gift from Dr. John Isaacs at The Johns Hopkins University. The rest was purchased from American Type Culture Collection (ATCC). Cells were cultured in RPMI or DMEM media with 5% FBS or charcoal-stripped FBS (CSS) and 1% penicillin streptomycin. Synthetic androgen R1881 was added at final concentration of 1 nM as androgen supplements in control conditions if needed. Enzalutamide and NAD+ inhibitors (OT-82 and FK866) were purchased from Selleckchem.

### Viability / proliferation

As previously reported ^39–41^, cell viability and proliferation experiments were carried out in 96 or 24 -well format, viable cells were determined with SYTO 60 fluoresce staining (Thermo Fisher), quantitated by LiCor Odyssey Imager, and expressed as % of solvent (DMSO, < 0.01% of final concentration) treated control.

### Single-cell RNA-Sequencing

Anti-androgen treatment naïve LNCaP cells (passage 5 from purchase) were treated with enzalutamide or solvent control for 24 hours. Afterwards, 100,000 cells from each condition underwent 10X chromium based single-cell RNA sequencing (10X Genomics) at OHSU massive parallel sequencing facility. Sequencing data was analyzed by Seurat packages (Seurat 4.0).

### Single cell-based colony formation and isolation

Anti-androgen treatment naïve LNCaP cells (passage 5 from purchase) were plated in single-cell suspension for ∼14 days to form single-cell based colonies, which were then isolated with pipette tips, resuspended and allowed to grow to single-colony derived clones in 96-well plates with androgen-containing media.

### NAPRT knockdown and overexpression

The siRNA cocktails containing 4-5 constructs against NAPRT were purchased from Sigma. Cells were transfected with silencing siRNA or non-targeting control siRNA for 24 hours, and then treated with NAD+ biosynthesis inhibitors to measure viability and proliferation. The lentiviral NAPRT overexpression plasmid was purchased from VectorBuilder. The transduction was based on procedures that were previously described ^42^. After the siRNA or overexpression, cells were treated with OT-82. In parallel experiments, cell lysates were harvested for western blot analysis to ensure NAPRT knockdown or overexpression.

### GFP and RFP transfection and fluorescence-based cell imaging / counting

As previously described ^42^, AR^high^ and AR^low/-^ cells were transduced with pseudo-lentivirus, containing plasmids coding for eGFP and RFP, respectively. Cells with stable GPF or RFP expressions were selected by puromycin. For fluorescence-based cell imaging in response to ADT/Enz, AR-high/GFP+ and AR-low/RFP+ cells were mixed and resuspended in R1881-containing media and plated into 96-or 24-well plates. After cells were attached overnight, cells were treated by control (CSS-media + 1 nM R1881) or ADT/Enz (CSS-media + 5 uM of enzalutamide). The images of GFP and RFP cells were taken with EVOS FL Auto Imaging System (Thermo Fisher Scientific). For proliferation analysis, GFP/RFP-transfected cells were plated in 96-well plate and cultured with incubators equipped with live camera (Incucyte Live Cell Analysis System). Cell pictures were taken every 4 hours for up to 96 hours. Cell viabilities were quantified based on total cell numbers or GFP/RFP intensities using the associated software.

### Western blotting and qRT-PCR

The antibodies (AR, Vimentin, NAPRT, NAMPT, Tubulin, β-actin) for western blots were purchased from Santa Cruze. All qRT-PCR primers were purchased from Integrated DNA Technologies (IDT). The western blot and qRT-PCR procedures were described in detail previously ^39–41^. Western blot was carried out with equal amount of whole cell lysates and visualized with LiCor Odyssey Fluorescence Imager. Tubulin or beta-actin were used as the equal loading control. The qRT-PCR was carried out with TRIzol-extracted RNA and SYBR-Green-based detection. The expression of beta-actin was used as the control.

### NAD+ measurements and rescue

The colorimetric kits for cellular NAD+ measurements were purchased from Abcam. The measurements were carried out based on manufacture’s instruction. For the rescue experiments, nicotinamide mononucleotide (NMN), the NAMPT product and NAD+ precursor, was purchased from Sigma, resuspended in cell culture media at the final concentration of 1 mM.

### Xenograft experiments

All animal experiments are in compliance with protocols approved by OHSU IACUC. As previously described^12^, subcutaneous implants of LNCaP clones were generated in male nude mice by injecting ∼5 million cells mixed with Matrigel. The dose for enzalutamide was 10 mg/kg (gavage), 5 times per week. The tumor volume was determined by digital caliper measurement, and expressed either as mm^3^ (volume = width x depth x length x 0.52) or as % of growth relative to the start of treatment.

### Statistical analysis

All experimental data were expressed as mean & standard deviation (s.d.) unless indicated otherwise. Statistical comparisons between two sample sets were performed with two-sided student’s *t*-test or paired *t*-test, comparisons among more than two samples were performed with repeated measures ANOVA, using MedCalc software. P < 0.05 was considered as significant, and P ≥ 0.05 was considered as not significant different (ND).

### Data availability

The sequencing data generated with this study is deposited in Gene Expression Omnibus (GEO) database under the accession code GSE120343. All data are available to the public. https://www.ncbi.nlm.nih.gov/geo/query/acc.cgi?acc=GSE120343. The prostate cancer TCGA and SU2C dataset referenced in the study are available in a public repository from the cBioPortal website (http://www.cbioportal.org/). The authors declare that all other data supporting the findings of this study are available within the article and its supplementary information files and from the corresponding author upon reasonable request.

## Author Contributions

DZQ designed all the experiments with contributions from Hao Geng, Hyun-Kyung Ko, Changhui Xue, and Kasen Shi. DZQ wrote the paper.

**The authors declare no competing interests.**

## Acknowledgment

We thank Drs. Gordan Mills and Dong Zhang for the use of microscopes and Incucyte. This work is supported by OHSU Prostate Cancer Program and NIH grants 5R01CA149253 and 5R01CA207377 to DZQ.

## Notes

### Competing Interest Statement

The authors have declared no competing interest.

## References

1. Denmeade SR, Isaacs JT. A history of prostate cancer treatment. Nat Rev Cancer. May 2002;2(5):389–396.

2. Watson PA, Arora VK, Sawyers CL. Emerging mechanisms of resistance to androgen receptor inhibitors in prostate cancer. Nat Rev Cancer. Dec 2015;15(12):701–711.

3. Tran C, Ouk S, Clegg NJ, et al. Development of a second-generation antiandrogen for treatment of advanced prostate cancer. Science. May 8 2009;324(5928):787–790.

4. Beer TM, Armstrong AJ, Rathkopf DE, et al. Enzalutamide in Metastatic Prostate Cancer before Chemotherapy. N Engl J Med. Jun 1 2014.

5. Bluemn EG, Nelson PS. The androgen/androgen receptor axis in prostate cancer. Curr Opin Oncol. May 2012;24(3):251–257.

6. Mills IG. Maintaining and reprogramming genomic androgen receptor activity in prostate cancer. Nat Rev Cancer. Mar 2014;14(3):187–198.

7. Davies AH, Beltran H, Zoubeidi A. Cellular plasticity and the neuroendocrine phenotype in prostate cancer. Nat Rev Urol. May 2018;15(5):271–286.

8. Brady SW, McQuerry JA, Qiao Y, et al. Combating subclonal evolution of resistant cancer phenotypes. Nat Commun. Nov 1 2017;8(1):1231.

9. Hinohara K, Polyak K. Intratumoral Heterogeneity: More Than Just Mutations. Trends Cell Biol. Jul 2019;29(7):569–579.

10. Lee MC, Lopez-Diaz FJ, Khan SY, et al. Single-cell analyses of transcriptional heterogeneity during drug tolerance transition in cancer cells by RNA sequencing. Proc Natl Acad Sci U S A. Nov 4 2014;111(44):E4726–4735.

11. Haffner MC, Zwart W, Roudier MP, et al. Genomic and phenotypic heterogeneity in prostate cancer. Nat Rev Urol. Feb 2021;18(2):79–92.

12. Geng H, Xue C, Mendonca J, et al. Interplay between hypoxia and androgen controls a metabolic switch conferring resistance to androgen/AR-targeted therapy. Nat Commun. Nov 26 2018;9(1):4972.

13. Urbanucci A, Barfeld SJ, Kytola V, et al. Androgen Receptor Deregulation Drives Bromodomain-Mediated Chromatin Alterations in Prostate Cancer. Cell Rep. Jun 06 2017;19(10):2045–2059.

14. Alumkal JJ, Sun D, Lu E, et al. Transcriptional profiling identifies an androgen receptor activity-low, stemness program associated with enzalutamide resistance. Proc Natl Acad Sci U S A. Jun 2 2020;117(22):12315–12323.

15. Asangani IA, Dommeti VL, Wang X, et al. Therapeutic targeting of BET bromodomain proteins in castration-resistant prostate cancer. Nature. Jun 12 2014;510(7504):278–282.

16. Kim DH, Sun D, Storck WK, et al. BET Bromodomain Inhibition Blocks an AR-Repressed, E2F1-Activated Treatment-Emergent Neuroendocrine Prostate Cancer Lineage Plasticity Program. Clin Cancer Res. Sep 1;27(17):4923–4936.

17. Kumar A, Coleman I, Morrissey C, et al. Substantial interindividual and limited intraindividual genomic diversity among tumors from men with metastatic prostate cancer. Nat Med. Apr 2016;22(4):369–378.

18. Nguyen HG, Yang JC, Kung HJ, et al. Targeting autophagy overcomes Enzalutamide resistance in castration-resistant prostate cancer cells and improves therapeutic response in a xenograft model. Oncogene. Sep 4;33(36):4521–4530.

19. Nouri M, Ratther E, Stylianou N, Nelson CC, Hollier BG, Williams ED. Androgen-targeted therapy-induced epithelial mesenchymal plasticity and neuroendocrine transdifferentiation in prostate cancer: an opportunity for intervention. Front Oncol. 2014;4:370.

20. Qi W, Morales C, Cooke LS, Johnson B, Somer B, Mahadevan D. Reciprocal feedback inhibition of the androgen receptor and PI3K as a novel therapy for castrate-sensitive and - resistant prostate cancer. Oncotarget. Dec 8 2015;6(39):41976–41987.

21. Kregel S, Chen JL, Tom W, et al. Acquired resistance to the second-generation androgen receptor antagonist enzalutamide in castration-resistant prostate cancer. Oncotarget. May 3 2016;7(18):26259–26274.

22. Bishop JL, Thaper D, Vahid S, et al. The Master Neural Transcription Factor BRN2 Is an Androgen Receptor-Suppressed Driver of Neuroendocrine Differentiation in Prostate Cancer. Cancer Discov. Jan 2017;7(1):54–71.

23. Rycaj K, Cho EJ, Liu X, et al. Longitudinal tracking of subpopulation dynamics and molecular changes during LNCaP cell castration and identification of inhibitors that could target the PSA-/lo castration-resistant cells. Oncotarget. Mar 22 2016;7(12):14220–14240.

24. Qin J, Liu X, Laffin B, et al. The PSA(-/lo) prostate cancer cell population harbors self-renewing long-term tumor-propagating cells that resist castration. Cell Stem Cell. May 4 2012;10(5):556–569.

25. Pfeiffer MJ, Schalken JA. Stem cell characteristics in prostate cancer cell lines. Eur Urol. Feb 2010;57(2):246–254.

26. Beltran H, Oromendia C, Danila DC, et al. A Phase II Trial of the Aurora Kinase A Inhibitor Alisertib for Patients with Castration-resistant and Neuroendocrine Prostate Cancer: Efficacy and Biomarkers. Clin Cancer Res. Jan 1 2019;25(1):43–51.

27. Coleman DJ, Gao L, King CJ, et al. BET bromodomain inhibition blocks the function of a critical AR-independent master regulator network in lethal prostate cancer. Oncogene. Apr 17 2019.

28. Venkadakrishnan VB, Presser AG, Singh R, et al. Lineage-specific canonical and non-canonical activity of EZH2 in advanced prostate cancer subtypes. Nat Commun. Aug 8 2024;15(1):6779.

29. Smith R, Liu M, Liby T, et al. Enzalutamide response in a panel of prostate cancer cell lines reveals a role for glucocorticoid receptor in enzalutamide resistant disease. Sci Rep. Dec 10;10(1):21750.

30. Mutetwa T, Foulkes WD, Polak P. Olaparib for Metastatic Castration-Resistant Prostate Cancer. N Engl J Med. Aug 27 2020;383(9):890.

31. Martinez-Pena F, Pearson AD, Tang EL, et al. Synthesis and biological evaluation of novel FiVe1 derivatives as potent and selective agents for the treatment of mesenchymal cancers. Eur J Med Chem. Nov 15 2022;242:114638.

32. Covarrubias AJ, Perrone R, Grozio A, Verdin E. NAD(+) metabolism and its roles in cellular processes during ageing. Nat Rev Mol Cell Biol. Feb 2021;22(2):119–141.

33. Zapata-Perez R, Wanders RJA, van Karnebeek CDM, Houtkooper RH. NAD(+) homeostasis in human health and disease. EMBO Mol Med. Jul 7 2021;13(7):e13943.

34. Lee J, Kim H, Lee JE, et al. Selective Cytotoxicity of the NAMPT Inhibitor FK866 Toward Gastric Cancer Cells With Markers of the Epithelial-Mesenchymal Transition, Due to Loss of NAPRT. Gastroenterology. Sep 2018;155(3):799–814 e713.

35. Piacente F, Caffa I, Ravera S, et al. Nicotinic Acid Phosphoribosyltransferase Regulates Cancer Cell Metabolism, Susceptibility to NAMPT Inhibitors, and DNA Repair. Cancer Res. Jul 15 2017;77(14):3857–3869.

36. Galli U, Colombo G, Travelli C, Tron GC, Genazzani AA, Grolla AA. Recent Advances in NAMPT Inhibitors: A Novel Immunotherapic Strategy. Front Pharmacol. 2020;11:656.

37. Korotchkina L, Kazyulkin D, Komarov PG, et al. OT-82, a novel anticancer drug candidate that targets the strong dependence of hematological malignancies on NAD biosynthesis. Leukemia. Jul 2020;34(7):1828–1839.

38. Bishop JL, Davies A, Ketola K, Zoubeidi A. Regulation of tumor cell plasticity by the androgen receptor in prostate cancer. Endocr Relat Cancer. Jun 2015;22(3):R165–182.

39. Geng H, Ko HK, Pittsenbarger J, et al. HIF1 and ID1 Interplay Confers Adaptive Survival to HIF1alpha-Inhibition. Front Cell Dev Biol. 2021;9:724059.

40. Liu Q, Geng H, Xue C, Beer TM, Qian DZ. Functional regulation of hypoxia inducible factor-1alpha by SET9 lysine methyltransferase. Biochim Biophys Acta. May 2015;1853(5):881–891.

41. Liu Q, Harvey CT, Geng H, et al. Malate dehydrogenase 2 confers docetaxel resistance via regulations of JNK signaling and oxidative metabolism. Prostate. Jul 2013;73(10):1028–1037.

42. Geng H, Rademacher BL, Pittsenbarger J, et al. ID1 enhances docetaxel cytotoxicity in prostate cancer cells through inhibition of p21. Cancer Res. Apr 15 2010;70(8):3239–3248.

